# Modification and Analysis of Context-Specific Genome-Scale Metabolic Models: Methane-Utilizing Microbial Chassis as a Case Study

**DOI:** 10.1101/2024.09.16.613288

**Authors:** M.A. Kulyashov, R. Hamilton, Y. Afshin, S.K. Kolmykov, T.S. Sokolova, T.M. Khlebodarova, M.G. Kalyuzhnaya, I.R. Akberdin

## Abstract

Context-specific genome-scale model (CS-GSM) reconstruction is becoming an efficient strategy for integrating and cross-comparing experimental multi-scale data to explore the relationship between cellular genotypes, facilitating fundamental or applied research discoveries. However, the application of CS modeling for non-conventional microbes is still challenging. Here, we present a GUI interface that integrates COBRApy, EscherPy, and RIPTiDe Python-based tools within the BioUML platform and streamlines the reconstruction and interrogation of the CS genome-scale metabolic frameworks via Jupyter Notebook. The approach was tested using -omics data collected for *Methylotuvimicrobium alcaliphilum* 20Z^R^, a prominent microbial chassis for methane capturing and valorization. We optimized the previously reconstructed whole genome-scale metabolic network by adjusting the flux distribution using gene expression data. The outputs of the automatically reconstructed CS metabolic network were comparable to manually optimized *i*IA409 models for Ca-growth conditions. However, the CS model questions the reversibility of the phosphoketolase pathway and suggested higher flux via primary oxidation pathways. The model also highlighted unresolved carbon partitioning between assimilatory and catabolic pathways at the formaldehyde-formate node. Only a very few genes and only one enzyme with a predicted function in C1-metabolism, a homolog of the formaldehyde oxidation enzyme (*fae1-2)*, showed a significant change in expression in La-growth conditions. The CS-GSM predictions agreed with the experimental measurements under the assumption that the Fae1-2 is a part of tetrahydrofolate-linked pathway. The cellular roles of the tungsten (W)-dependent formate dehydrogenase (*fdhAB*) and *fae*-homologues (*fae1-2,* and *fae3*) were investigated via mutagenesis. The phenotype of the f*dhAB* mutant followed the model prediction. Furthermore, more significant reduction of the biomass yield was observed during growth in La-supplemented media, confirming a higher flux through formate. *M. alcaliphilum* 20Z^R^ mutants lacking *fae1-2* did not display any significant defects in methane or methanol-dependent growth. However, contrary to *fae,* the *fae1-2* homolog failed to restore the formaldehyde activating enzyme function in complementation tests.

Overall, the presented data suggest that the developed computational workflow supports the reconstruction and validation of CS-GSM networks of non-model microbes.

## Introduction

Complete and accurate reconstruction of metabolic networks is central for advancing fundamental discoveries or rebuilding microbial systems toward desired outcomes (Orth et al., 2010; Bordbar et al., 2014; Sahu et al., 2021). Model reconstruction includes numerous steps, but in general success relies on the efficient integration of multi-scale experimental data, such as structural and functional organization of genomes, transcriptomes, proteomes, and metabolomes. The integration requires additional mathematical model developments that can capture changes in intra- and extracellular metabolites of a cell in response to genetic and/or environmental perturbation (Esvelt and Wang, 2013). The model is expected to provide *in silico* and yet reliable solutions that reproduce the systems behaviors such as growth rate or cell yields, and predict genetic alterations required to improve desired outcomes (i.e., product yield) under defined growth conditions (Gu et al., 2019; Walakira et al., 2021).

Advances in next-generation sequencing approaches and automatic reconstruction of genome-scale metabolic (GSM) models have led to active application of network-guided metabolic engineering of non-model organisms, often with unique metabolisms like methane utilization and lithotrophy. Numerous models are now available in BiGG (Norsigian et al., 2020), BioModels (Malik-Sheriff et al., 2020) and MetaFishNet (Li et al., 2010). GSM reconstruction can further facilitate developments in non-model microbial systems via the addition of omics-driven constraints on the distribution of fluxes to realistically represent metabolic capabilities of the organism of interest (Zhang and Hua, 2016; Aite et al., 2018).

The present study describes the computational framework for semi-automated reconstruction of a context-specific (CS) model of non-model organisms. The workflow includes a set of Python programs including COBRApy (Ebrahim et al., 2013), EscherPy (King et al., 2015) and RIPTiDe (Jenior et al., 2020) integrated into the Jupyter Notebook environment on the BioUML platform (Kolpakov et al., 2022). The pipeline was tested for the CS-GSM reconstruction of methane-utilizing bacteria (known as methanotrophs^)^. There is a significant interest in compiling the unique metabolic capabilities of methanotrophic bacteria for biotechnological and/or environmental applications. We anticipate that the computational framework presented below can expedite the implementation of GSM models for such advancements. We selected *Methylotuvimicrobium alcaliphilum* 20Z^R^, as over years the *Methylotuvimicrobium spp* has become a testable microbial platform for methane capturing and valorization (Nguyen et al., 2019; Awasthi et al., 2022; Villada et al., 2022; He et al., 2023; Ngoc Pham et al., 2023). The metabolic flux balance models of the whole genome have been generated and manually curated (Torre et al., 2015; Akberdin et al., 2018a; Stone et al., 2020), allowing for a direct comparison between expert-curated and automated optimization of GSMs for non-model microbes.

## 1 Materials and Methods

### 1.1 Strain, growth media and -omics datasets

#### Strains

The lab strain of *Methylotuvimicrobium alcaliphilum* 20Z^R^ is a gram-negative, gamma proteobacterium that can grow on either methane or methanol as its main source of carbon (Khmelenina et al., 1999). The cultures of *Methylobacterium extorquence* AM1 and *Methylobacterium extorquence* AM1 Δ*fae* were kindly provided by Dr. N. Cecilia Martinez-Gomez (UC Berkeley) (also see Table 3). The following *Escherichia coli* strains were used for cloning and gene transfer experiments: *E.coli* S17-1 (lab stock), and *E.coli* DH10B (NEB Labs, Catalog Number C3019H: *Δ(ara-leu) 7697 araD139 fhuA ΔlacX74 galK16 galE15 e14-ϕ80dlacZΔM15 recA1 relA1 endA1 nupG rpsL (Str^R^) rph spoT1 Δ(mrr-hsdRMS-mcrBC)*).

#### Growth media

M. *alcaliphilum* 20Z^R^ strains were routinely grown in P_3%_ nitrate mineral media supplemented with methanol (0.2% v/v) or methane as described previously (Collins and Kalyuzhnaya, 2018). Methane (25-50 ml) was introduced into 125-250 ml vials with 25-50 ml media using syringe to keep a methane:air ratio of 20:80 in headspace. Plates were incubated in anaerobic jars and refilled using an Anoxomat system programmed to provide an atmosphere of 12-25% methane and 75-88% air. *M. extorquence* strains were grown using Hypho media (Harder et al., 1973), supplemented with methanol (0.2% v/v) or succinate (0.4% v/v). Liquid cultures were incubated at 30°C with shaking at 200 rpm. Plates were kept in 30°C incubators. *E.coli* strains were grown using LB broth or LB agar media (Miller, Difco BD Life Sciences) and incubated at 37°C. The following antibiotics were applied when required: kanamycin (100µg/ml), rifamycin (50µg/ml).

#### Gene expression studies

The cultivation parameters of M. *alcaliphilum* 20Z^R^, culture growth rates and sequencing details were previously described (Akberdin et al., 2018a) and are summarized in Table 1. Briefly, the cultivation parameters and cell samples for RNA sequencing were collected from steady-state bioreactor cultures growing in nitrate mineral media at pH 8.7-9 with or without La-supplementation. The parameters for each bioreactor unit (250 ml vessel with 125-150 ml of media) were set with agitation at 500 rpm, temperature kept at 30°C, and gas inflow rate of 0.3 sL/hr.

**Table 1.** Summary of RNA-seq transcriptomic datasets used for the optimization of the GSM model.

| Sample name | Sample id | Raw reads | Aligned reads | Dataset ID | CH<sub>4</sub>/O<sub>2</sub> Consumption | Growth rate, hr<sup>-1</sup> |
| --- | --- | --- | --- | --- | --- | --- |
| **Ca-BR1** | Ca-CH4-BR1_S1_L005_R1_001 | 22124770 | 10 140 552 | GSM8020389 | 1.12 ± 0.09 | 0.05 |
| **CaBR2** | Ca-CH4-BR2_S1_L005_R1_001 | 17884237 | 16 955 446 | GSM8020390 | 1.12 ± 0.09 | 0.05 |
| **LaBR1** | La-CH4-BR1_S1_L005_R1_001 | 18542763 | 8 409 524 | GSM8020393 | 1.28 ± 0.01 | 0.07 |
| **LaBR2** | La-CH4-BR2_S1_L005_R1_001 | 16381413 | 12 029 600 | GSM8020394 | 1.28 ± 0.01 | 0.07 |

The RNA-seq reads were mapped to the *M. alcaliphilum* 20Z^R^ reference genome (ASM96853v1) using the Bowtie2 algorithm v2.5.1 (Langmead and Salzberg, 2013). Illumina standard adapters were removed using Trim Galore (Kreuger, 2021) software (when necessary) before mapping. The mapped reads were quantified with FeatureCounts (Liao et al., 2014) using gene features from the RefSeq annotation (GCF_000968535.2). To extend the functional annotation of mapped genes, a comparison of three annotations (old and new RefSeq as well as Genoscope annotations) for this genome was carried out (Supplementary Table 1). Subsequently, DESeq2 (Love et al., 2017) was used to normalize and perform differential expression analysis. Genes are considered to be differentially expressed if they have |log2(fold change)| > log2(1.5) at adjusted p-value < 0.05. The EnhancedVolcano R-package was used to visualize the obtained results.

For KEGG signaling pathways enrichment analysis we used a homebrew R-script. Specifically, the KEGGREST R-package was applied to match genes to KEGG signaling pathways. Then Wilcoxon rank-sum test was performed for enrichment testing. The analysis was performed independently on either the full set of genes or specifically on genes considered in gene-protein-reaction associations of the model.

The results of the analyses are stored on the BioUML platform (Kolpakov et al., 2022) and available by the following link: https://gitlab.sirius-web.org/RSF/20ZR_CS_GSM_model. The RNA-seq data (counts and differentially expressed genes) were submitted to the NCBI Gene Expression Omnibus (GEO) database under accession number <u>GSE253414</u>.

### 1.2 Extension of the *i*IA409 model, Flux Balance Analysis with COBRApy and model visualization with EscherPy

To integrate transcriptomic data into the mathematical model, a published and experimentally verified mathematical model for 20Z^R^, *i*IA409 (Akberdin et al., 2018a) was harnessed. The model was modified to account for the presence of metals either in the medium or transported into the cell that were not previously considered explicitly. In addition, different scenarios for the formaldehyde partitioning between tetrahydrofolate (H_4_F) and tetrahydromethanopterin (H4MPT) pathways and phosphoketolase (EC4.1.2.22 and EC 4.1.2.9) were evaluated and compared with the original model. Modifications of the original (i.e., *i*IA409) model as well as a subsequent flux balance analysis of the modified *i*IA409-based models and CS models reconstructed using the RIPTiDe algorithm (Jenior et al., 2020), were made via the COBRApy library (v. 0.25.0; Ebrahim et al., 2013) using Jupyter widgets. The program code is given at the link (https://gitlab.sirius-web.org/RSF/20ZR_CS_GSM_model/-/tree/master/Data/CS_GSM_model/file_collection.files).

The EscherPy library (v. 1.7.3; King et al., 2015) was employed to visualize flux distributions predicted by the modified and CS models. The final network is available at the link (https://gitlab.sirius-web.org/RSF/20ZR_CS_GSM_model/-/tree/master/Data/CS_GSM_model/file_collection.files).

The differential reaction fluxes (DRFs) between two model predictions were visualized on the metabolic map if the ratio of fluxes for a reaction fell within the boundaries of 0.67 <DRF <1.5 (i.e., the change in the reaction flux occurs more than 1.5 times). Additionally, reactions that were activated (On) or turned off (Off) in the CS model but not in the original, were assigned to a ratio value of 1000 (for On) or 2000 (for Off). To visually distinguish these turned on/off reactions from other DRFs, a specific color assignment was applied: pink for the value of 1000 (On reactions) and aqua for the value of 2000 (Off reactions). The visualization program code is given under the link in the Jupyter Notebook called “20Z_transcript” in sections: “Escher visualization” and “DRF analysis” (https://gitlab.sirius-web.org/RSF/20ZR_CS_GSM_model/-/tree/master/Data/CS_GSM_model/file_collection.files).

### 1.3 Integration of transcriptomics data into the model via RIPTiDe

The RIPTiDe algorithm (v.3.4.79; Jenior et al., 2020) was used to integrate transcriptomic data. Counts reflecting expression levels for each gene in TPMs (transcript per million) in the corresponding growth conditions (Table 1) were used as a source of transcriptomic data for the construction of CS models. Right before data integration, the selected changes were made to the model to match the selected cultivation conditions, and, if necessary, the biomass metal content was modified using the original code implemented within the Jupyter Notebook using the Jupyter widgets modules, which greatly simplified the process of the model modification. The stoichiometric coefficients for La^3+^ and W^4+^ in the biomass equation were set at experimentally measured concentrations (summarized in Table 2). Also, the transcriptomic data were prepared using the RIPTiDe algorithm. Because we used data normalized by DESeq2, the normalization step with RIPTiDe was skipped. The file for the algorithm was converted to TSV format, containing a column with gene IDs (the same as those used in gene-protein-reaction (GPR) rules in the *i*IA409 model) and columns with normalized counts for each gene. For direct reconstruction of CS models, the *maxfit* module was used, which analyzes models with different proportions of the original biomass in the model and selects the one in which the predicted fluxes distribution had the highest level of correlation with transcriptomic data (Table 2). RIPTiDe *contextualize* module was applied to reanalyze steps which were not calculated by original *maxfit.* All codes for the model modification and transcriptomic data integration are provided within the BioUML platform and available at https://gitlab.sirius-web.org/RSF/20ZR_CS_GSM_model/-/tree/master/Data/CS_GSM_model/file_collection.files.

**Table 2.**
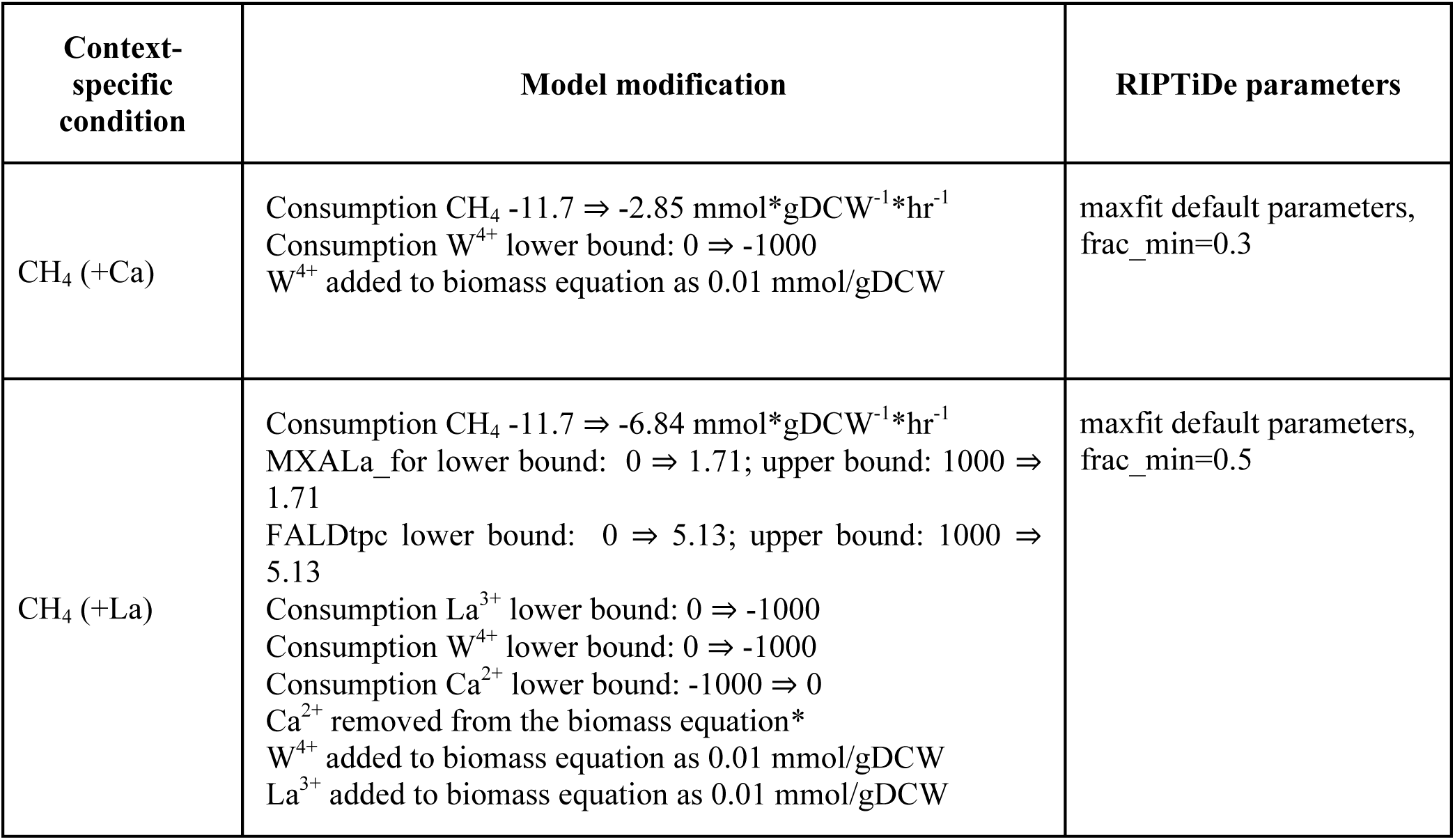
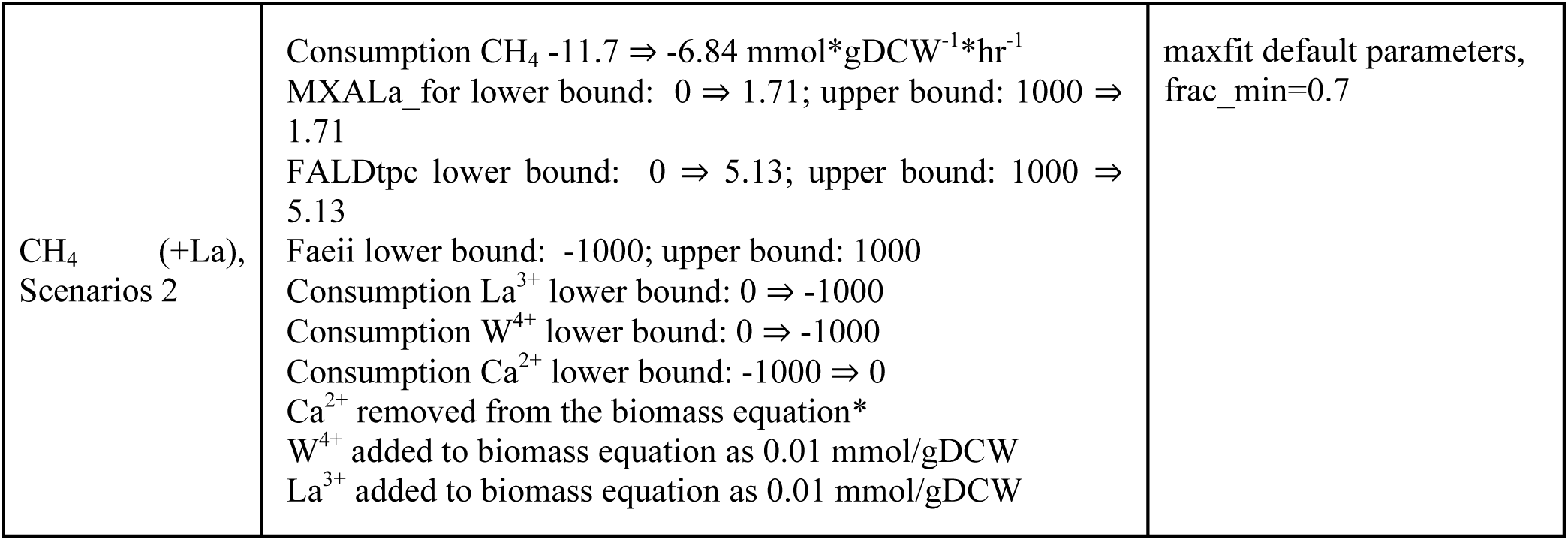
Modifications and *in silico* growth conditions used for the reconstruction of CS models via RIPTiDe algorithm.

| Context-specific condition | Model modification | RIPTiDe parameters |
| --- | --- | --- |
| $\text{CH}_4$ (+Ca) | Consumption $\text{CH}_4$ -11.7 ⇒ -2.85 mmol*gDCW<sup>-1</sup>*hr<sup>-1</sup><br>Consumption $\text{W}^{4+}$ lower bound: 0 ⇒ -1000<br>$\text{W}^{4+}$ added to biomass equation as 0.01 mmol/gDCW | maxfit default parameters, frac_min=0.3 |
| $\text{CH}_4$ (+La) | Consumption $\text{CH}_4$ -11.7 ⇒ -6.84 mmol*gDCW<sup>-1</sup>*hr<sup>-1</sup><br>MXALa_for lower bound: 0 ⇒ 1.71; upper bound: 1000 ⇒ 1.71<br>FALDtpc lower bound: 0 ⇒ 5.13; upper bound: 1000 ⇒ 5.13<br>Consumption $\text{La}^{3+}$ lower bound: 0 ⇒ -1000<br>Consumption $\text{W}^{4+}$ lower bound: 0 ⇒ -1000<br>Consumption $\text{Ca}^{2+}$ lower bound: -1000 ⇒ 0<br>$\text{Ca}^{2+}$ removed from the biomass equation*<br>$\text{W}^{4+}$ added to biomass equation as 0.01 mmol/gDCW<br>$\text{La}^{3+}$ added to biomass equation as 0.01 mmol/gDCW | maxfit default parameters, frac_min=0.5 |
| CH<sub>4</sub> (+La),<br>Scenarios 2 | Consumption CH<sub>4</sub> -11.7 ⇒ -6.84 mmol*gDCW<sup>-1</sup>*hr<sup>-1</sup><br>MXALa_for lower bound: 0 ⇒ 1.71; upper bound: 1000 ⇒ 1.71<br>FALDtpc lower bound: 0 ⇒ 5.13; upper bound: 1000 ⇒ 5.13<br>Faeii lower bound: -1000; upper bound: 1000<br>Consumption La<sup>3+</sup> lower bound: 0 ⇒ -1000<br>Consumption W<sup>4+</sup> lower bound: 0 ⇒ -1000<br>Consumption Ca<sup>2+</sup> lower bound: -1000 ⇒ 0<br>Ca<sup>2+</sup> removed from the biomass equation*<br>W<sup>4+</sup> added to biomass equation as 0.01 mmol/gDCW<br>La<sup>3+</sup> added to biomass equation as 0.01 mmol/gDCW | maxfit default parameters,<br>frac_min=0.7 |

### 1.4 Fae phylogeny, construction of mutant strains and phenotyping

#### Phylogeny

More than 560 formaldehyde activating enzymes (Fae) and or Fae-like proteins with some homology to the Fae*-*protein from *M. alcaliphilum* 20Z^R^ (Gene ID: 2540616361; MEALZ_RS11875) were identified in the Uniprot database. Of the 560 sequences, 78 were selected for phylogenetic comparison. The sequences represent 39 species—3 archaea and 36 bacteria, including planctomycetes, firmicutes, actinobacteria and proteobacteria and a representative from the NC10 phylum. The amino acid sequence of archaeal *fae*-homologue (*Geoglobus ahangarhi*, Gene ID: 2810347888; Ga0325130_111788) was selected as the outgroup, as it had the lowest bootstrap value at 0.19% indicating it as the most distant Fae sequence. Alignments were completed using the MAFT v7 E-INS algorithm (Katoh and Standley, 2013). MEGAX (Tamura et al., 2021) was used for best predicted model LG+G and 1000 bootstrap replicates were performed when constructing the tree.

#### Complementation tests

The schematic for plasmid construction and integration of 20Z^R^ *fae* into *M. extorquens* AM1Δ*fae* is presented in Supplementary Figure 1. To construct *fae*-expression vectors, the *fae*-genes from 20Z^R^ and pAWP78(P_89_) plasmid were amplified by PCR using Flash Phusion master mix (Thermo Fischer) using *M. alcaliphilum* DNA. The *fae* genes were then integrated into the pAWP78 vector under P_89_ promoter by Gibson assembly using NEBuilder (NEB Labs) and transformed into *E.coli* DH10B competent cells. Promoter P_89_ is a truncated hybrid version of the P_tac_ promoter, with a fragment of the P_trp_, *lac*-operator, and a mxaF ribosome binding site (AGGAAA). The promoter is known to provide constitutive expression in methylotrophic bacteria (Puri et al., 2015). The correct assemblies of the genetic constructs were validated by sequencing. The vectors were subcloned into *E.coli* S17-1. The plasmids were then transferred via conjugation into an *M. extorquens* AM1Δ*fae* mutant via bi-parental mating. pAWP78 empty vector (EV) was cloned into wild type *M. extorquens* AM1 and the mutant strain AM1Δ*fae* to construct control variants (Table 3). Lawns of *M. extorquens* AM1Δ*fae* supplemented with succinate (1-2 day growth) and lawns of pAWP78-P*_89_*-20Z^R^*fae* in *E.coli* S17-1 (overnight) were prepared for mating on plates with Hypho media supplemented with 5% Nutrient Broth (NB Difco, BD Life Sciences). Plates were incubated for 1 day at 30^°^ C. The biomass was collected and spread onto Hypho-agar plates supplemented with succinate and kanamycin for selection. Kanamycin-resistant clones were observed after 3-4 days of incubation. Clones were transferred into Hypho-agar plates with kanamycin and rifamycin for *E.coli* counter selection.

**Table 3.**
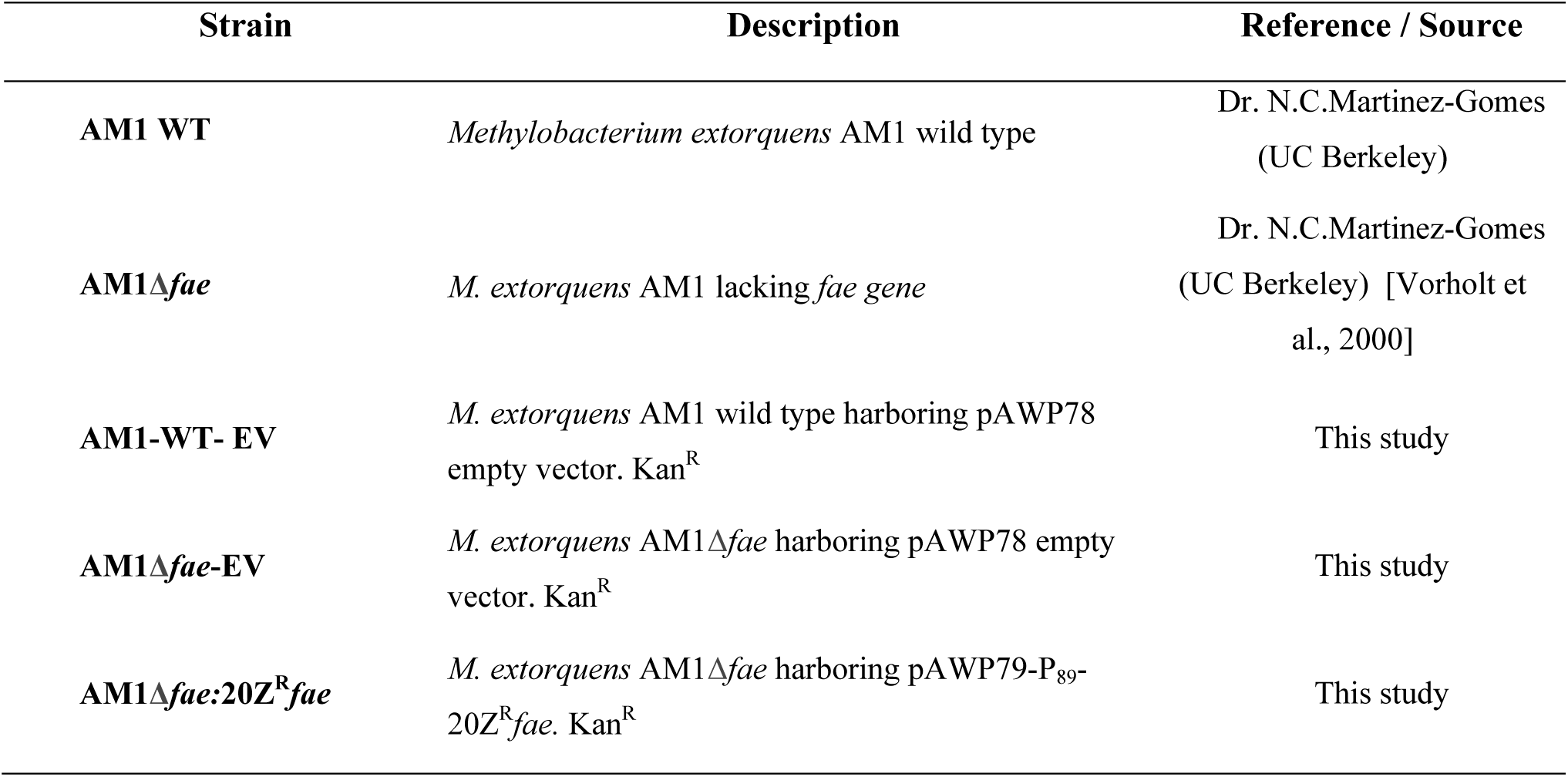
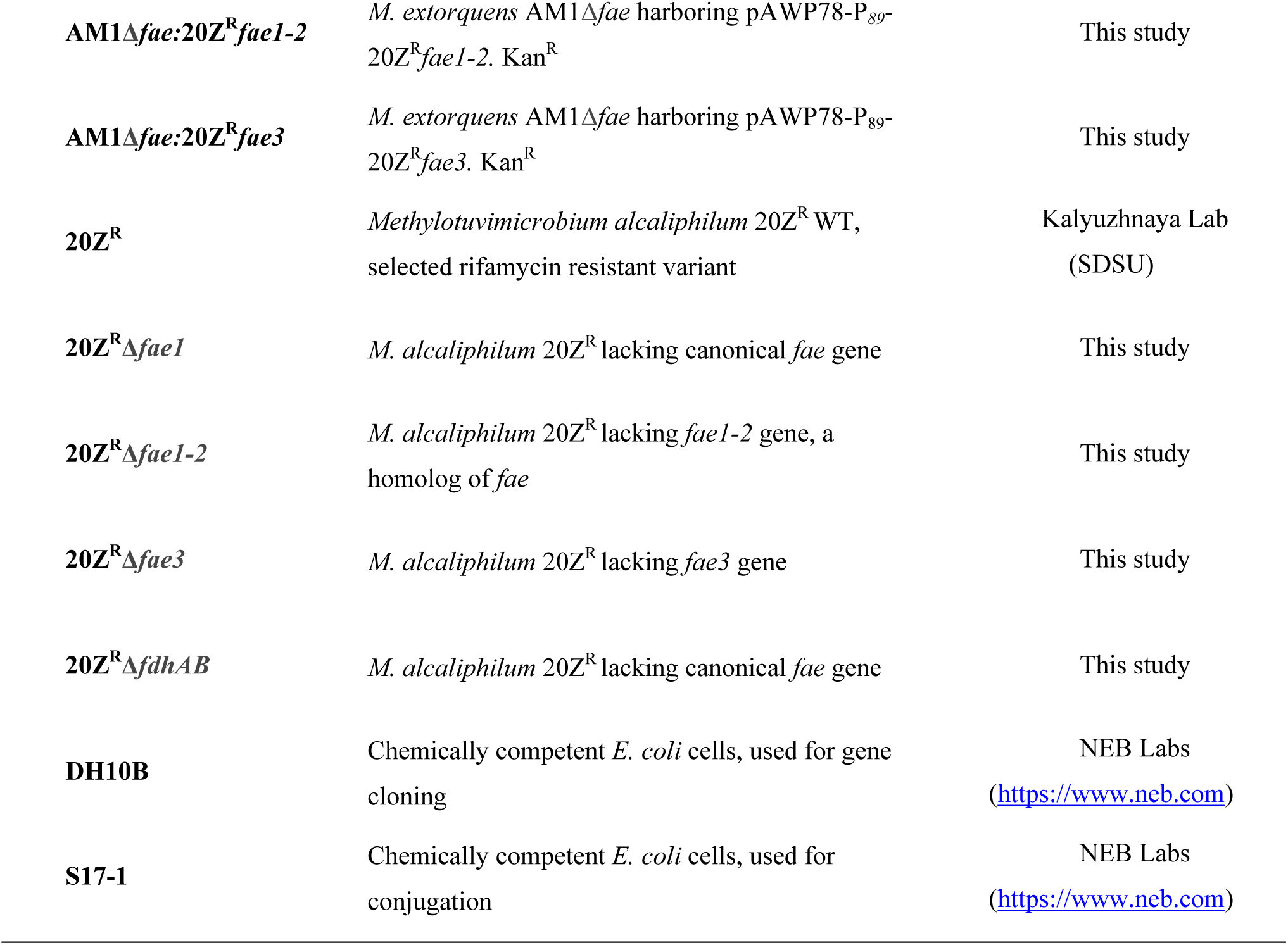
Strains used in this study.

The *M. extorquens* AM1Δ*fae* -mutant complementation tests were conducted using: 1. **Control:** Hypho medium supplemented with succinate (0.4%); 2. **Sensitivity test:** Hypho medium supplemented with succinate (0.4%) and methanol (0.2%); 3. **Growth test:** Hypho medium supplemented with methanol (0.2%). Cultures were incubated at 30^°^ C and evaluated after 1-2 days (succinate plates) and after 3-4 days (methanol or methanol + succinate plates).

Liquid batch culture experiments were carried out in flasks containing 25 mL of Hypho and supplemented with succinate (0.4%) and/or methanol (0.2%). At least three biological replicates with two technical replicates each per culture were tested. Each biological replicate was inoculated from a single colony. Growth measurements were taken starting from OD 0.1 until reaching the stationary phase and were used to calculate growth rates and standard deviations were calculated to produce graphic representation of data acquired.

#### Mutagenesis

All mutants were constructed using the allelic exchange vector pCM433 (Marx, 2008). Upstream and downstream flanking regions of *fae* genes or *fdhAB*-gene cluster (MEALZ_1882-MEALZ_1883) genes from *M. alcaliphilum* 20Z^R^ were amplified in using 2x Fusion Phlash (Thermo Fischer). The pCM433 vector was amplified in the same way using pCM433-F and pCM433-R primers (Collins and Kalyuzhnaya, 2018). The PCR products were run on a gel and purified. The flanking regions were integrated into pCM433 vector using a NEBuilder HiFi DNA assembly master mix. Reactions were transformed into *E. coli* DH10B competent cells. Kanamycin-resistant colonies were selected. Plasmids were purified using Zymoclean Gel DNA recovery kit (Zymo Research, USA). The constructs were verified by PCR and sequences were confirmed by sequencing. The resulted plasmids were transformed into *E.coli* S17-1 and mated with *M. alcaliphilum* 20Z^R^. Kanamycin-resistant colonies were subjected for negative selection on sucrose. The genotype of resulted clones was confirmed by PCR and PCR-fragment sequencing. All constructed strains are listed in Table 3.

## 2 Results

Below, we summarize the main steps of the reconstruction and analysis of a CS GSM (Figure 1) using the previously developed GSM model of *M. alcaliphilum* 20Z^R^ (Akberdin et al., 2018a) and the RIPTiDe algorithm (Jenior et al., 2020).

**Figure 1.**
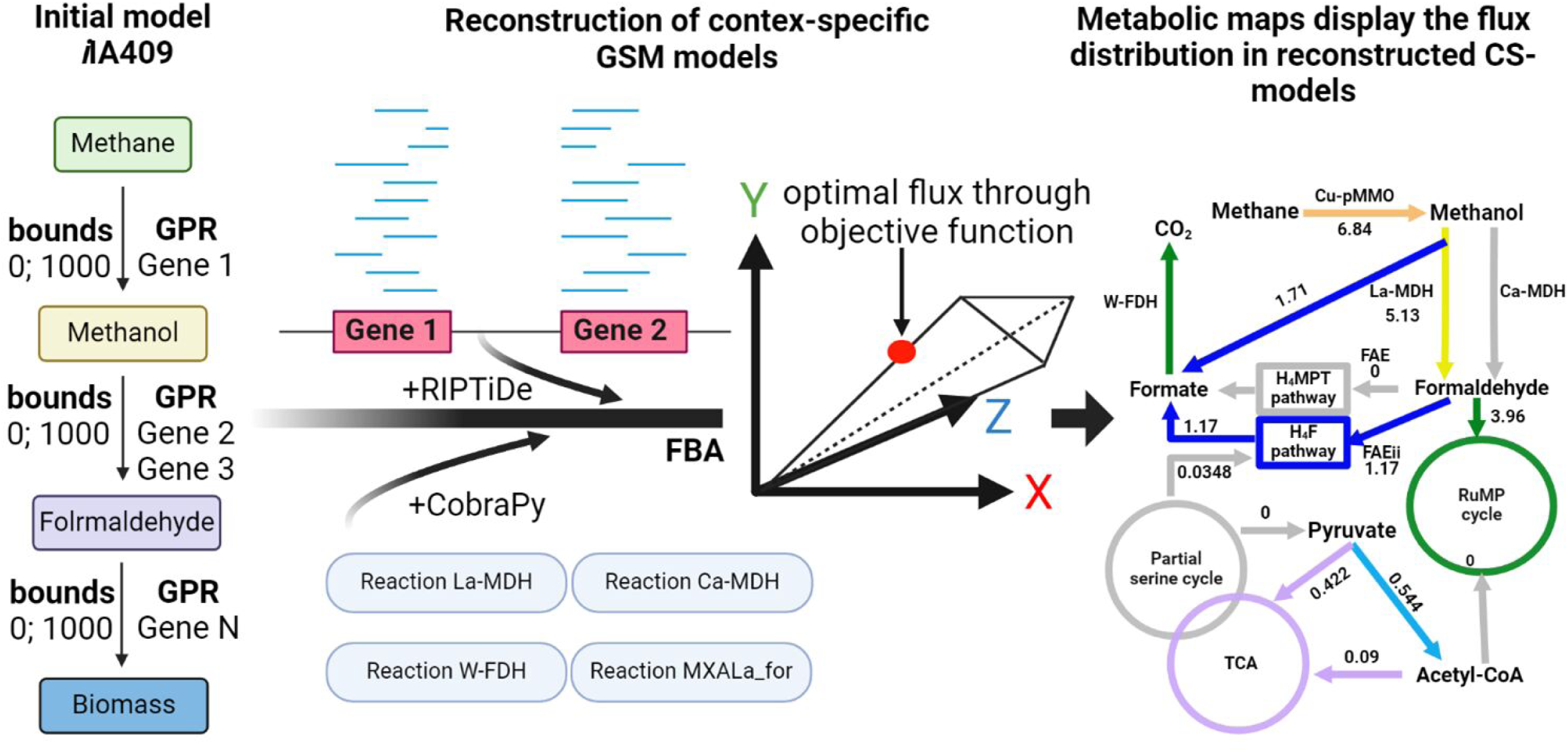
A schematic representation of the developed pipeline to reconstruct and analyze the CS GSM model for *M. alcaliphilum* 20Z^R^. At the first step, the original model was extended using COBRApy (Ebrahim et al., 2013) to account for the presence in the medium and transport into the cell of some metals (see Materials and Methods). Thereafter, the transcriptomics data for different growth conditions (or contexts) were integrated into the extended model to reconstruct a number of CS models using the RIPTiDe tool (Jenior et al., 2020). Finally, flux balance analysis (FBA) was conducted in COBRApy and predicted fluxes distributions were displayed on the metabolic maps using the Escher web-tool (King et al., 2015). All steps of the pipeline are performed in the corresponding Jupyter Notebook (see Availability Statement).

### 2.1 Construction of CS-GSM model for aerobic methanotroph *M. alcaliphilum* 20Z^R^

To build a workflow for the CS modeling with GSM and available -omics data, we applied Jupyter Notebook (Kluyver et al., 2016; Perkel, 2018). The computational platform included Python-based tools such as COBRApy (Ebrahim et al., 2013), EscherPy (King et al., 2015) and RIPTiDe (Jenior et al., 2020) to simplify the process of handling GSM models.

To reconstruct CS-GSMs for *M. alcaliphilum* 20Z^R^, the original GSM model was extended. Specifically, the Ca^2+^ transport reactions were added as an environmental control of the enzymatic reaction catalyzed by Ca^2+^-dependent methanol dehydrogenase (Ca-MDH, *mxaFI*) (Figure 2A). Similarly, the transport reaction for La^3+^ was added and linked with the La-dependent methanol dehydrogenase (La-MDH, *xoxF*) reaction (Figure 2B). Since it has been shown that La-MDHs can oxidize methanol to formate (Pol et al., 2014), the corresponding reaction was added in addition to the methanol to formaldehyde conversion. These modifications enabled tunable control of the methanol conversion steps depending on the cultivation conditions. This modification also allows control of the methanol flux via Ca-MDH or La-MDH accordingly to their gene expression levels. Additionally, a transport reaction for tungsten (W) was added and integrated with the reaction catalyzed by W-dependent formate dehydrogenase (Figure 2C).

**Figure 2.**
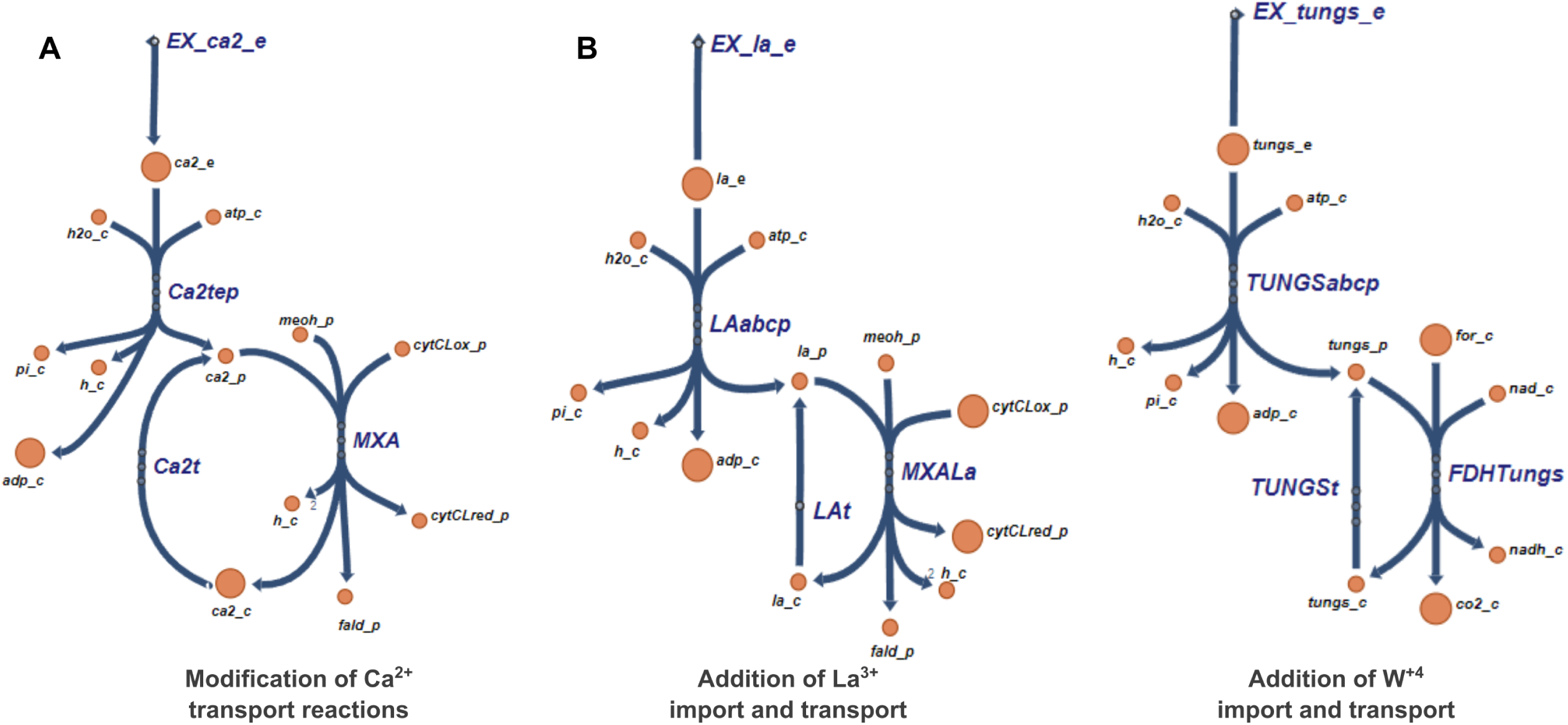
Visualization of the reactions that were added to the model for calcium (A), lanthanum (B) and tungsten (C) via Escher tool (King et al., 2015). A. Modified Ca^2+^ transport reactions; B. Added exchange and transport reactions for La^3+^; C. Added exchange and transport reactions for W^4+^.

The growth rates predicted by the *i*IA409 and optimized CS models were similar, but both below the experimentally observed for Ca-growth conditions (Supplementary Table 2). However, the predicted O_2_:CH_4_ ratios were similar to the experimental measurements (1.22 and 1.273 for *i*IA409 and CS model, respectively vs 1.12 ± 0.09 obtained for chemostat cultures).

Similarly to the original model, the predicted O_2_ consumption reaction rate was significantly higher than experimental measurements for the La-growth conditions too (see details in Supplementary Table 2). The introduction of O_2_ uptake constraints to the CS-model tuned the *in-silico* prediction toward measured parameters. Despite similar outputs, some differences between the two types of *M. alcaliphilum* genome-scale metabolic models were observed, in particular, differences in predicted growth rates, suggesting variations in carbon flux distributions. To investigate this further we carried out comparative analysis of the fluxes and applied network visualization (Escher tool) to highlight the most significant divergences.

### 2.2. CS-GMS highlights unresolved carbon distribution during growth in the presence of lanthanides

The Ca- and La-growth conditions were investigated separately. Detailed Escher metabolic maps are presented in Supplementary Figures 1.2-1.3 for Ca-growth condition and in Supplementary Figures 1.5-1.6 for La growth condition. The CS model predicts a slightly higher activity of the H4MPT pathway (from 1.07 to 1.25 mmol/gDCW/hr compared to the original model) and activation of the uphill electron transport mechanism unlike to the original model for Ca-growth (Supplementary Figures 1.2 - 1.4).

The flux distribution predicted by the CS model for growth with La differed more substantially from that predicted by the original model. To quantitatively identify metabolic differences between the original model and the CS model, the metabolic map was constructed using DRFs (see Methods). The constructed map for La growth conditions showed differences in value of fluxes of reactions related to the RuMP pathway: reactions associated with ribulose-5-phosphate regeneration are less active in the CS model simulation (decrease flux through transaldolase (TALA) and transketolase reactions (TKT1 and TKT2) in 1.37 times, from 1.28 to 0.933 and 1.31 to 0.952 mmol/gDCW/hr, respectively). Similar trend was observed for pentose-5-phosphate 3-epimerasereaction (PPE) from 2.59 to 1.88 mmol/gDCW/hr). Carbon flux via the he EMP and EDD pathways also decreased compared to the original model predictions in 1.37 times. The decreased flux of CO_2_ production reaction (in 1.67 times, from 3.55 to 2 mmol/gDCW/hr) was accompanied by lower CO_2_ production via the formate dehydrogenase (FDH) reaction in 1.9 times (from 2.95 to 1.56 mmol/gDCW/hr), which like the Ca-growth condition, however to the higher degree (Supplementary Figures 1.5 - 1.7).

At the same time, the carbon flow via the C1 transfer pathways was increased. The high flux via H_4_MTP pathway can be linked with a very high-level expression of *fae1-2,* a homologue of canonical formaldehyde-activating enzyme (FAE), also known as 5,6,7,8-tetrahydromethanopterin hydro-lyase (Vorholt et al., 2000). *fae1-2* expression was more than 4 times higher during La-supplemented growth, while the expression of other genes in the H_4_MPT pathway did not significantly change (Supplementary Figure 1.8, Table 4). Furthermore, a 2-fold reduction was observed for the canonical FAE. The differential expression of the *fae*-genes between Ca and La growth conditions has been previously noted, and while the alternative metabolic function for *fae1-2* has been speculated, it remains unresolved (Akberdin et al., 2018a).

**Table 4.** Comparison of transcription level for FAE genes (*fae1, fae1-2, fae3*) between Ca- and La-presents conditions.

| Gene ID | Number of<br>normalized counts<br>+Ca |  | Number of<br>normalized counts<br>+La |  | log2Fold<br>Change | P<sub>adj</sub> |
| --- | --- | --- | --- | --- | --- | --- |
| MEALZ_RS11875<br>(*fae1*) | 22233 | 21377 | 14086 | 13351 | -0.431 | 0.001 |
| MEALZ_RS04100<br>(*fae1-2*) | 2560 | 1834 | 40690 | 24419 | 4.094 | 7.01E<sup>-17</sup> |
| MEALZ_RS07105<br>(*fae3*) | 6392 | 5327 | 3519 | 3245 | -0.55 | 0.00026 |

To evaluate the *fae*-homologue’s contribution to carbon flux deterring from the RuMP pathway towards the primary oxidation pathway, we reconstructed the CS model which assumes different roles for the Fae1-2, from its contribution to Fae1 function to the carbon flow into the H_4_Folate pathway by adding the reaction FAEii which is the condensation reaction of formaldehyde with tetrahydrofolate and considering different ratios of formate and formaldehyde produced by La-MDH (*xoxF)* (Figure 3, Table 5).

**Figure 3.**
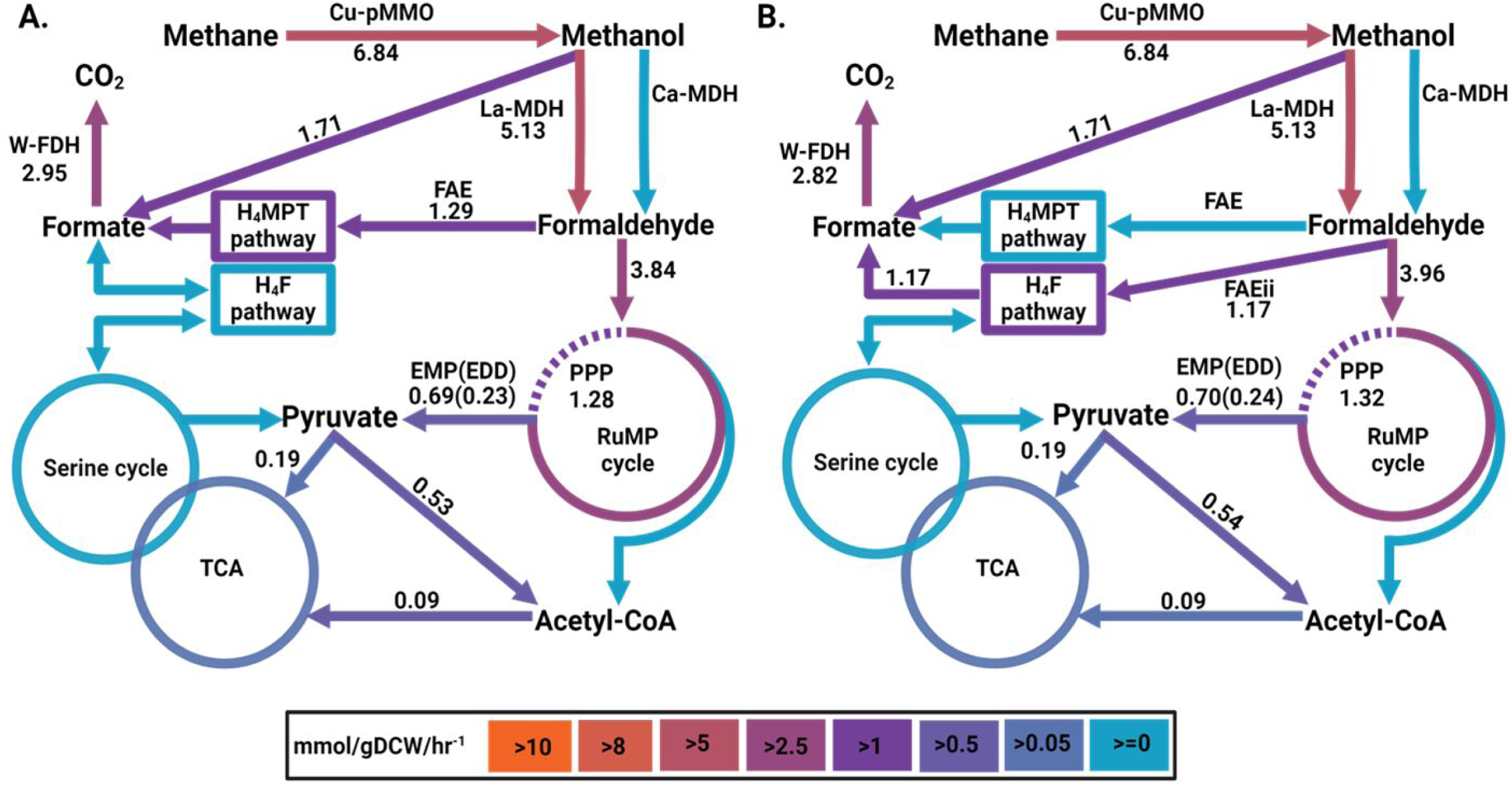
The schematic representation of the fluxes distribution predicted **(A)** by original *i*IA409 model for growth on CH_4_ in the presence of La, W, Cu with FAEii in H4MTP mode and **(B)** by CS model based on transcriptomic counts data for growth on CH_4_ in the presence of La, W, Cu with FAEii in H4Folate mode. The line thickness and color indicate the extent of the flux rate through the reaction in mmol*gDCW^-1^*hr^-1^. Abbreviations on the figure means: **Cu-pMMO:** a reaction catalyzed by Cu**-**dependent pMMO; **La-MDH** and **Ca-MDH:** reactions with activities of two MDH isoforms (La**-** and Ca**-**dependent respectively); **W-FDH:** the enzymatic reaction catalyzed by W-dependent FDH. **FAEii –** reaction which describes the potential role of *fae1-2* in condensation of formaldehyde and tetrahydrofolate.

**Table 5.**
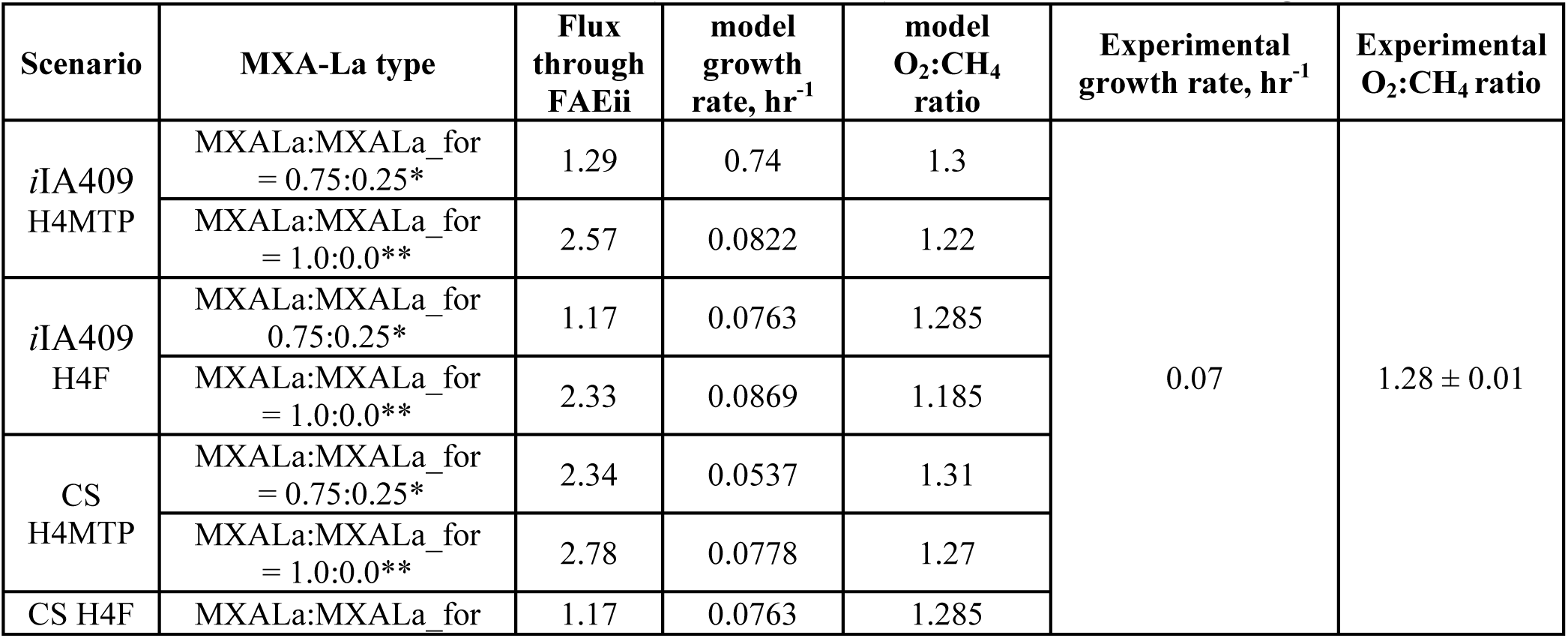
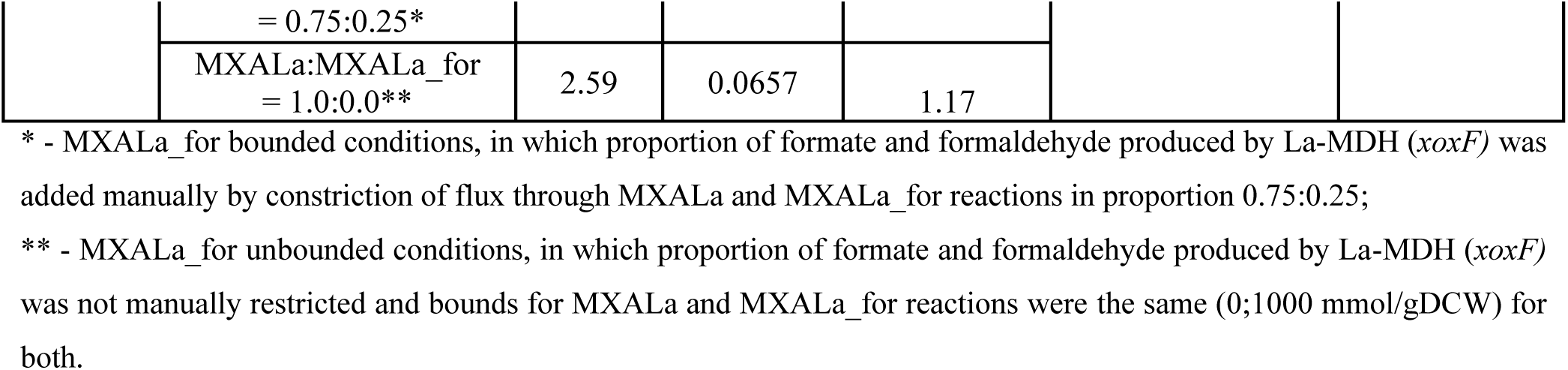
Predicted flux via the cofactor (H_4_F or H_4_MTP) mediated C_1_-transfer for growth with La.

We also investigated an opportunity of the reverse function for Fae1-2 (hydrolysis instead of condensation). The assumption of Fae1-2 role in the H_4_Folate pathway provided the best fit to the experimental data (Table 5) for both the original and CS models. Furthermore, the CS-model in this condition or the context predicts O_2_ consumption rate without any restrictions on its reactions unlike CS-model without FAEii reaction. Considering that La-growth contributes to a higher formate pool, the essentiality of the Fae1-2 function becomes more apparent. To further investigate the prediction, additional targeted mutagenesis studies were carried out. First, we investigated the impact of the formate dehydrogenase knockout on cell growth with Ca versus La. Then we investigated phenotypes of *fae1-2* knockouts. The functional characterization of Fae1-2 was supplemented by phylogenetic and complementation studies.

#### Mutagenesis

A knockout strain 20Z^R^ Δ*fdhAB* was constructed as described in Materials and Methods. The growth phenotypes of the mutant were tested using growth media containing either La or Ca to investigate the impacts of the La-switch (i.e., XoxF methanol dehydrogenase instead of Ca-dependent MxaFI). The 20Z^R^Δ*fdh* knockouts displayed growth defects when compared to the wild type (WT), and the growth reduction was more severe during La-supplementation (Figure 4). The growth rate was almost 5-fold lowed in the *fdhAB* mutant compared to the wild type. No significant growth defects were observed in media supplemented with methanol (data not shown), suggesting that the FDH is more critical for methane utilization.

**Figure 4.**
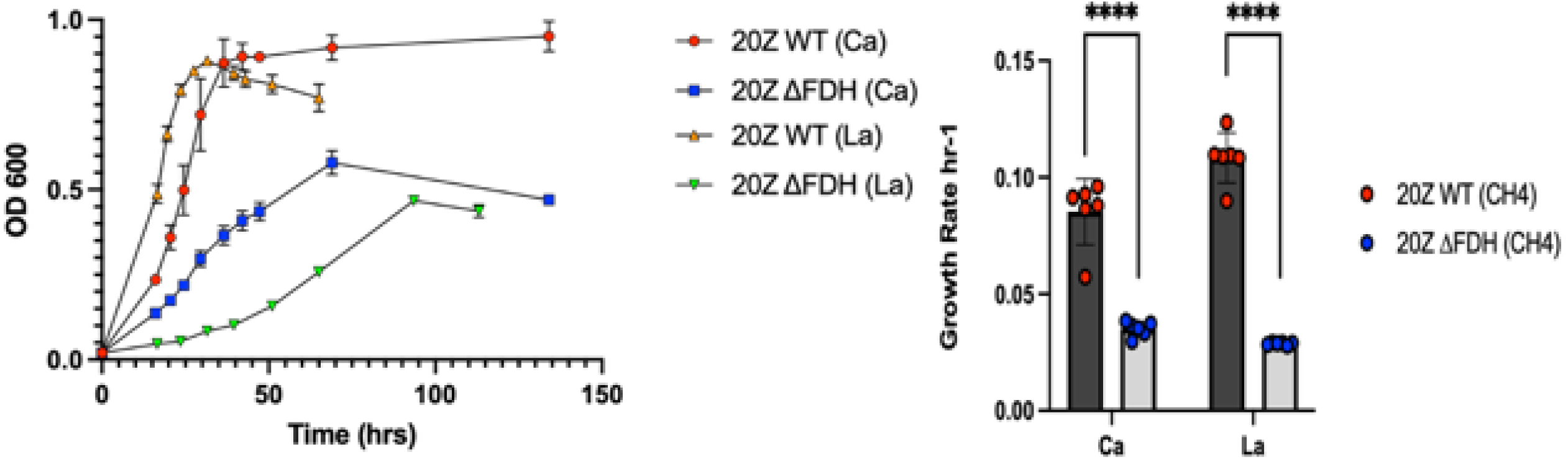
Growth parameters of *M. alcaliphilum* 20Z^R^ WT and 20Z^R^Δ*fdhAB* grown with methane using growth media supplemented with Ca or La.

We also constructed *M. alcaliphilum* mutants lacking *fae1-2*; however, the mutant did not display any growth defects (data not shown). We were not able to generate *fae1* knockouts, which indicates its essentiality for the growth.

#### Complementation tests

In order to investigate the functional activity of the Fae1 and Fae1-2 homologues, complementation studies were carried out. *M. extorquens* AM1Δ*fae,* the null mutant strain, does not have the ability to use methanol as a carbon source and is highly sensitive to methanol (Vorholt et al., 2000). The strain AM1 has been used for investigating the function of Fae-like sequences obtained from pure cultures as well as metagenomic sequences (Kalyuzhnaya et al., 2005; Crowther et al., 2008). In this study three plasmids, harboring *fae (*MEALZ_RS11875*), fae*1-2 (MEALZ_RS04100) and *fae3* (MEALZ_RS07105) homologues from *M. alcaliphilum* 20Z^R^ were constructed and integrated into *M. extroquens* AM1Δ*fae*. All strains have the ability to grow on succinate (Figure 5). When tested on solid media, both the sensitivity test and the growth test revealed that the integration of *fae* and *fae1-2* homologues into AM1Δ*fae* relieved methanol sensitivity through complementation. The strain AM1Δ*fae:*pAWP78-P*_89_*-20Z^R^*fae*3 displayed similar sensitivity to methanol as the parental AM1Δ*fae* strain, while both AM1Δ*fae:* pAWP78-P*_89_*-20Z^R^*fae* and AM1Δ*fae:*pAWP78-P*_89_*-20Z^R^*fae*1-2 become resistant to methanol. The *fae* and *fae1-2* genes also rescued the AM1Δ*fae* growth on methanol on solid media.

**Figure 5.**
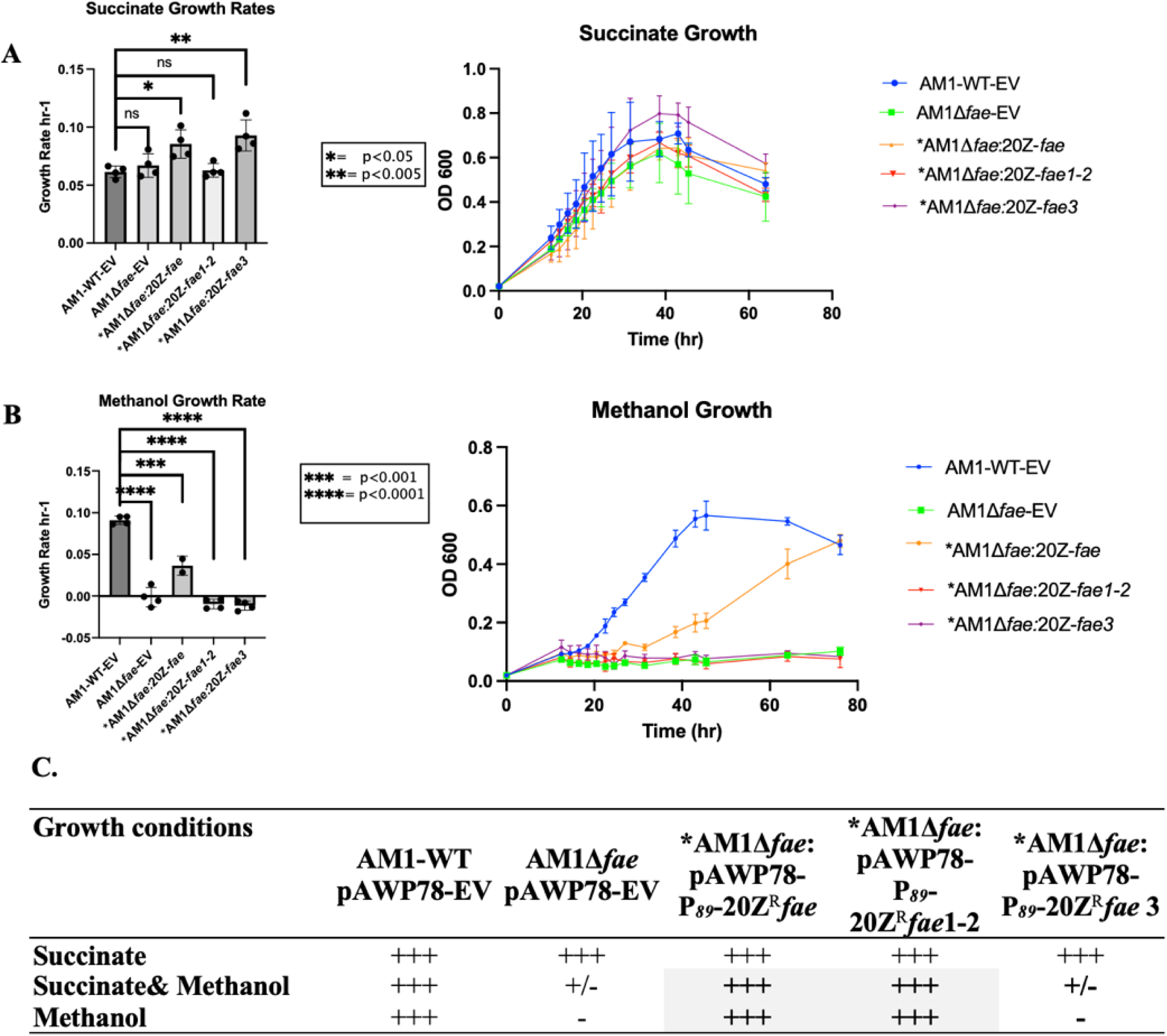
Growth parameters (growth rate, left and growth curve, right) of the *M. extorquens* AM1 wild type (WT) and AM1Δ*fae* strains expressing *fae* and *fae*-homologues on succinate (A) and methanol (B). **C.** Complementation results obtained on solid media. **\*** Complementation of the *AM1Δfae*-mutant growth on solid media with methanol is highlighted in grey.

The growth of the strains was also investigated in liquid culture and produced surprisingly different outcome for the Fae1-2 construct. While both, *fae1* and *fae1-2* restored growth on methanol on solid plates, only *fae1* is able to restore the AM1Δ*fae*-mutant phenotype. AM1Δ*fae*:20Z*fae1-2* is able to relieve methanol sensitivity on agar medium but is not able to grow in liquid culture supplemented with methanol (Figure 5B). AM1*Δfae,* AM1*Δfae:*20Z^R^*fae1-2* and AM1*Δfae:*20Z^R^*fae3* do not grow in liquid cultures supplemented with methanol.

#### Phylogenetic analyses of Fae-homologues

To analyze the functional roles of the *fae*-homologues and more specifically the La-induced *fae1-2* gene product, we carried out phylogenetic analyses. Fae and Fae-like proteins are homologous across different groups of bacteria such as the firmicutes, actinomycetes, planctomycetes proteobacteria and NC10 phylum, as well as archaea. The phylogenetic tree we constructed includes 78 *Fae* sequences that represent different microbial species, including 34 bacterial species and 5 archaeal. *Geoglobus ahangarhi* (Gene ID: 2810347888; Ga0325130_111788) roots the tree. Five main groups were identified. Only sequences from the group 1 (bacterial Fae) and group 5 (archaeal Fae) have defined cellular function (Figure 6). Group 1 Fae proteins capture formaldehyde generated from primary C_1_-oxidation (methanol to formaldehyde). Group 5 Fae-enzymes contribute to capturing formaldehyde produced by a reverse hexulose phosphate synthase activity during conversion of C6-sugars into C5-sugars for nucleotide biosynthesis (Goenrich et al., 2005; Grochowski et al., 2005). Both Group 1 and Group 5 Faes catalyze the formation of methylene-H_4_MPT from H_4_MPT and formaldehyde and thus serves as the entry point for the formaldehyde oxidation pathway. The percent identity shared for archaeal sequences in group 5 with canonical Fae from *M. extorquens* AM1 (Gene ID: 644811916; MexAM1_META1p1766) is between 50-51%.

**Figure 6.**
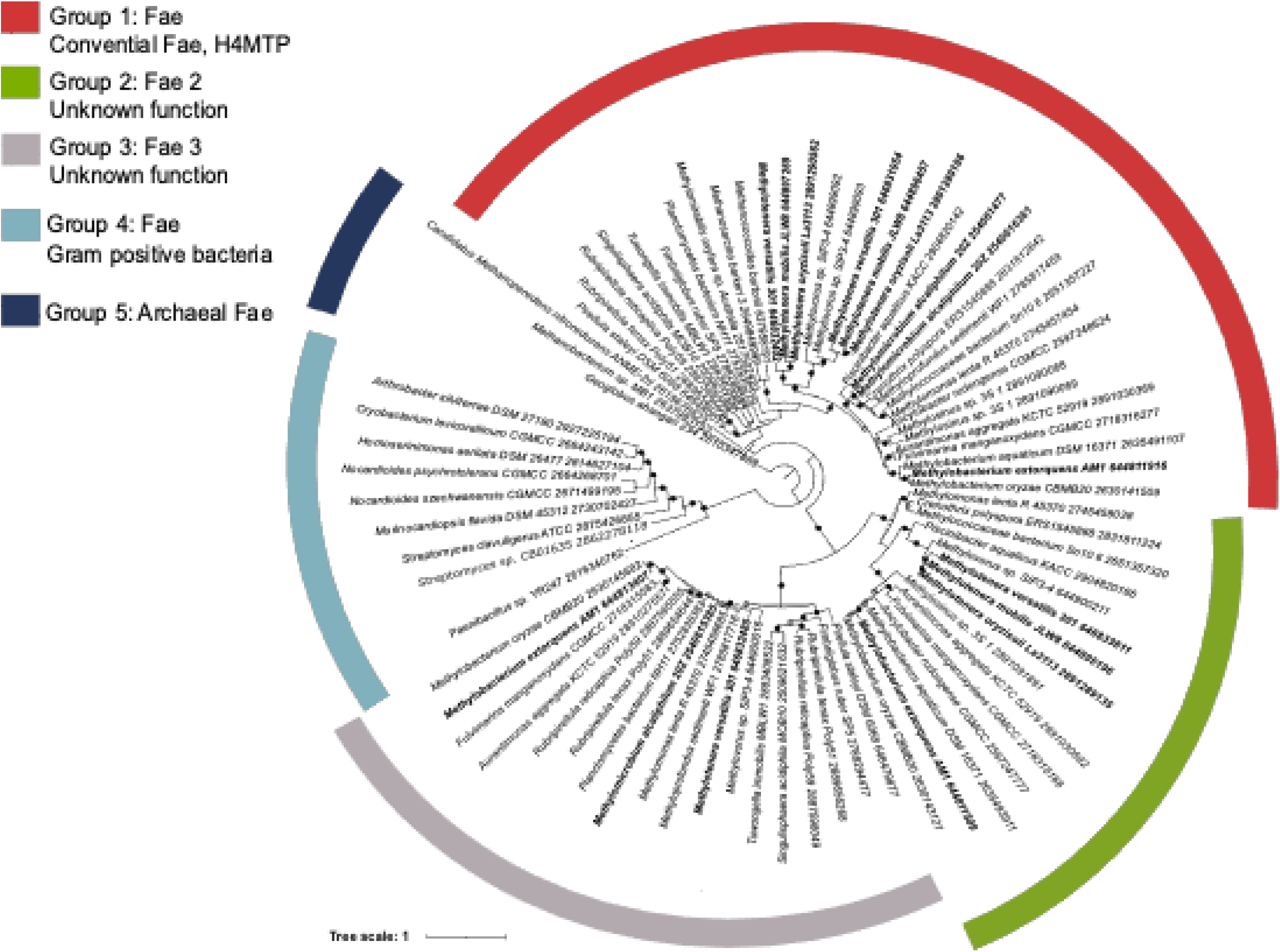
Phylogenetic analysis of Fae-homologues in bacteria and archaea. Protein sequences of Fae from *M. alcaliphilum* 20Z^R^, *Methylotenera* spp, and *M. extorquens* AM1 are highlighted in bold. Filled circles on nodes represent >70% bootstrap support; open circles on nodes represent >50% bootstrap support.

The cellular function of Fae-like proteins from Groups 2, 3 and 4 are not known. Suggested functions include regulatory or sensing roles or a possible contribution to tetrahydrofolate-linked C-transfer (Chistoserdova, 2011; Good et al., 2015).

## 3 Discussion and Conclusions

In this study we implemented the Python-based pipeline to reconstruct the context specific metabolic network of *M.alcaliphilum* 20Z^R^, a model methanotrophic bacterium. The initial goal of the study was to streamline metabolic network reconstruction for non-model microbial systems with unique metabolic capabilities, i.e., methanotrophy. The model reconstruction was followed up by the flux balance analysis for 20Z^R^ growing on methane with Ca or La supplementation. Considering that a great deal of information is available for the train, the reconstructed model was compared to previously expert-curated reconstructions (*i*IA409 model, Akberdin et al., 2018a).

The CS-model predictions were comparable to the original *i*IA409 model simulation with a few exceptions. The CS model indicated that reverse electron transfer through the cytochrome bc1 complex is essential for supporting methane oxidation, whereas the original model suggested a more direct redox input from a reduced cytochrome cL, a product of methanol oxidation. Therefore, the CS model predicted the necessity of the reverse electron transfer systems (ETS) without requiring additional expert-curated constraints, which were necessary for the original model (Akberdin et al., 2018b). It should be mentioned however, that both models, do not support the redox model of methane oxidation. The CS-model also ruled out a possibility of carbon flux from acetyl-P toward pentose phosphate pathway intermediates via the phosphoketolase pathway. Finally, CS-model suggested higher carbon flux via formaldehyde-formate nodes. The direct conversion to formate due to XoxF activity and the H_4_MPT pathway activation led to the enhanced production of formate compared to the value predicted by the original model. The increase of the expression of the gene encoding FDH has been confirmed by the analysis of DEGs between La and Ca conditions (Johnson et al., 2021). Follow-up studies with the formate dehydrogenase knockouts further confirm the essentiality of the formate oxidation reaction.

The CS-model predictions for the La-dependent growth condition demonstrated a set of metabolic rearrangements in comparison to the flux distributions obtained by simulation of the original model. The key difference in the carbon flux distribution predicted by the CS versus the original models is the increased role of the C1-transfer pathway. It has been shown previously that La-supplementation results in the reduction of gene expression for a canonical *fae,* while induces its close homolog, named here as *fae1-2*. We interrogated the possible functional role of *fae1-2* using the CS model. The model highlights a possibility that Fae1-2 is an enzyme that contributes to formaldehyde condensation with H_4_folate. Genetic tests further indicate that the Fae1-2 function differs substantially from the canonical Fae.

In summary, we demonstrate that this pipeline can help reconstruct metabolic models that are similar to manually curated networks. The pipeline for CS model development implemented in the BioUML platform can be used and easily tuned for the investigation of metabolic changes in other growth conditions of the methanotrophic strain. Furthermore, the reconstructed models should be able to highlight previously overlooked pathways, thus advancing fundamental knowledge of non-model microbial systems and promoting their development toward biotechnological or environmental implementations.

## 6 Author Contributions

IRA and MGK designed and coordinated the study. MAK reconstructed the CS GSM model for *M. alcaliphilum* 20Z^R^ and conducted FBA analysis of the updated model. RH carried out fermentation, RNA-seq studies and *fdhAB* mutagenesis. YA carried out *fae* mutagenesis and complementation studies. TSK and SKK conducted RNA-seq analysis. MAK, TMK, IRA and MGK analyzed the data and FBA results. MAK, SKK, TMK, IRA and MGK wrote the manuscript. All authors read and approved the final manuscript.

## 8 Supplementary Material

**Supplementary Table 1.** The functional annotation of mapped genes and a comparison of three annotations (old and new RefSeq as well as Genoscope annotations) for 20ZR genome.

**Supplementary Table 2.** Simulation results of the original *i*IA409 model and CS-GSMs for different growth conditions.

**Supplementary Figures 1.1-1.8.** Caption for each Supplementary Figure is presented in the file.

## 9 Data Availability Statement

The RNA-seq data (counts and differentially expressed genes) is available via GEO database under accession number GSE253414 (https://www.ncbi.nlm.nih.gov/geo/query/acc.cgi?acc=GSE253414). The described pipeline for transcriptomics data analysis, results of the DEGs analysis as well as the CS GSM model for 20ZR with the corresponding Jupyter Notebook are available on Gitlab at https://gitlab.sirius-web.org/RSF/20ZR_CS_GSM_model accessed on 14th August 2024.

